# Bayesian modeling and simulation to inform rare disease drug development early decision-making: application to Duchenne muscular dystrophy

**DOI:** 10.1101/2021.02.05.429907

**Authors:** Janelle L. Lennie, John T. Mondick, Marc R. Gastonguay

## Abstract

Rare disease clinical trials are constrained to small sample sizes and may lack placebo-control, leading to challenges in drug development. This paper proposes a Bayesian model-based framework for early go/no-go decision making in rare disease drug development, using Duchenne muscular dystrophy (DMD) as an example. Early go/no-go decisions were based on projections of long-term functional outcomes from a Bayesian model-based analysis of short-term trial data informed by prior knowledge based on 6MWT natural history literature data in DMD patients. Frequentist hypothesis tests were also applied as a reference analysis method. A number of combinations of hypothetical trial designs, drug effects and cohort comparison methods were assessed.The proposed Bayesian model-based framework was superior to the frequentist method for making go/no-go decisions across all trial designs and cohort comparison methods in DMD. The average decision accuracy rates across all trial designs for the Bayesian and frequentist analysis methods were 45.8 and 8.98%, respectively. A decision accuracy rate of at least 50% was achieved for 42 and 7% of the trial designs under the Bayesian and frequentist analysis methods, respectively. The frequentist method was limited to the short-term trial data only, while the Bayesian methods were informed with both the short-term data and prior information. The specific results of the DMD case study were limited due to incomplete specification of individual-specific covariates in the natural history literature data and should be reevaluated using a full natural history dataset. These limitations aside, the framework presented provides a proof of concept for the utility of Bayesian model-based methods for decision making in rare disease trials.

## Introduction

### Rare disease drug development challenges

Available treatments exist for only 5% of rare diseases, leaving hundreds of millions of affected patients worldwide without treatment [1]. Though advances have been made in recent years, treatments are needed for most of the 7,000 known rare diseases [1]. Key challenges to rare disease drug development include but are not limited to low numbers of patients, poor understanding of disease pathology and progression, lack of established clinical trial endpoints and surrogate biomarkers, and high variability in disease progression and presentation [2]. Small populations lead to poorly powered trials with small sample sizes; between 2006-2014, less than 50 subjects were recruited in 71.4% of registered rare disease trials [3]. Rare disease trials may lack placebo-control due to the ethical responsibility to treat patients suffering from life-threatening serious conditions, half of whom are children [1]. Standard of care or active treatments may serve as the reference arm in clinical trials, but often there is no effective standard of care. Clinical trials in rare diseases are necessarily constrained by small sample sizes, which also makes parallel control arms a challenge.

Typical drug development programs for common diseases include a Phase 2 proof-of-concept clinical trial to gain a preliminary understanding of potential clinical efficacy before investing in larger and more complex dose-ranging and confirmatory clinical trials. While the same objective is relevant to rare disease drug development, performing multiple, sequential trials is often not feasible. In many cases, only two trials are conducted for a rare disease: a combined Phase 1/2 trial to learn about efficacy and safety, and a combined Phase 2/3 trial to confirm efficacy. The trials are typically small and poorly powered for a clinical outcome endpoint and dose ranging trials are often limited to the objectives of safety, tolerability, and pharmacokinetics.. A central issue with drug approvals in rare diseases is the limited amount of evidence collected in these small clinical trials.

Despite these challenges, the regulatory standards for approval of rare disease drugs are similar to those applied to investigational treatments of common diseases; that is the requirement of demonstrating “substantial evidence” of efficacy and safety for the intended indication from adequate and well-controlled investigations [**?**, 4].

### Early decision-making strategies for rare disease drug development

Early decision-making strategies for drug development aim to increase the efficiency of compound selection for late stage development. Analyses of preliminary short-term studies are performed, yielding predictions of the probability of success in longer, larger scale clinical trials. Informed decisions to either continue (“go”) or terminate (“no-go”) development of the drug for the specific indication are made. Various statistical methods and decision criteria or stopping guidelines have been applied to go/no-go decisions [5–7].

### Bayesian model-based methods for drug development

The Bayesian statistical paradigm is a formal approach to include information that has been learned and accumulated prior to performing a new experiment. Prior knowledge is combined with newly collected data, and inferences are based on posterior probabilities of outcomes or quantities of interest.

When applied to drug development, Bayesian methods formally incorporate prior knowledge about the disease, drug, endpoints, or other factors with newly collected clinical trial data. Prior data may be borrowed from sources such as historical controls, disease consortia, patient registries, or previous trials. The totality of evidence is applied to robustly evaluate drug effects, bolstering statistical power. This is an appealing factor for rare disease drug development where limited evidence from small trials is a key challenge.

Bayesian inferences may be posed in terms of posterior probability distributions of model parameters or other quantities of interest, and predictive probability distributions of future events. Using Bayesian methods for drug development yields inferences based on clinically meaningful quantities specific to the indication. The probabilities are clearly and directly interpreted to assess trial results. Inference based on probabilities of meaningful clinical outcomes is appealing compared to hypothesis tests based on arbitrary levels of statistical significance.

### Duchenne muscular dystrophy drug development

Duchenne muscular dystrophy (DMD) is a genetic pediatric rare disease that has recently faced drug development challenges. The disease is characterized by a lack of endogenous functional dystrophin, which causes progressive muscle weakening and deterioration [8]. The 6-minute walk test (6MWT) has been a routinely utilized primary clinical endpoint for DMD efficacy trials [9, 10], with a 30-meter placebo-corrected change from baseline (ΔΔ6MWT) minimal clinically important difference (MCID) [11]. Recent DMD trials have targeted demonstration of the 30-meter MCID in trials ranging between 24-48 weeks long [**?**, 12].

The first approved drug for DMD was granted conditional accelerated approval in 2016 [13]. The initial attempt to demonstrate clinical benefit based on the primary functional endpoint was not achieved in a study of 12 patients [12, 14]. Instead, the accelerated approval was based on the effect of treatment on the surrogate biomarker, dystrophin, and was contingent upon verification of clinical benefits via the functional outcome in future long-term confirmatory trials [14]. This set a precedent for DMD drug development, and two more drugs were later granted the same type of conditional accelerated approval from the results of studies with 12 and 16 total subjects [15, 16].

Hypotheses based on other muscular dystrophies and animal models postulate that recovery of at least 10 [14] or 20 [17] percent of normal dystrophin in a DMD patient would likely demonstrate a clinical functional benefit. However, the relationship is mostly unclear and the amount of dystrophin “reasonably likely” to produce a clinical benefit has not been determined [17]. The percent of normal dystrophin recovered after treatment with each of the three approved drugs varied widely from a mean range of 0.93 - 5.7%, with none reaching the 10-20% range [14–16]. The challenge remains to achieve DMD drug approval based on a clinically meaningful functional endpoint.

### Objectives

This paper proposes a Bayesian model-based framework for early go/no-go decision making in rare disease drug development, using DMD as an example. Early go/no-go decisions are based on projections of long-term functional outcomes from a Bayesian model-based analysis of simulated short-term trial data informed by prior knowledge based on 6MWT natural history data in DMD patients. Posterior simulations provide predictions of long-term 6MWT distributions to inform early decision making in the context of a clinically relevant assessment time frame. A number of hypothetical trial designs and analysis methods are assessed. The overall goal of this paper is to illustrate an example of early decisions made from projections of long-term outcomes, informed by prior natural history data. The resulting framework could be applied to any rare disease drug development program.

## Methods

### Overview

The methods for the model-based analysis followed a simulation-estimation-simulation archetype. A natural history model was augmented with a hypothetical drug effect model to use as a clinical trial simulation and estimation model in this work. (Fig 1)

**Fig 1.**
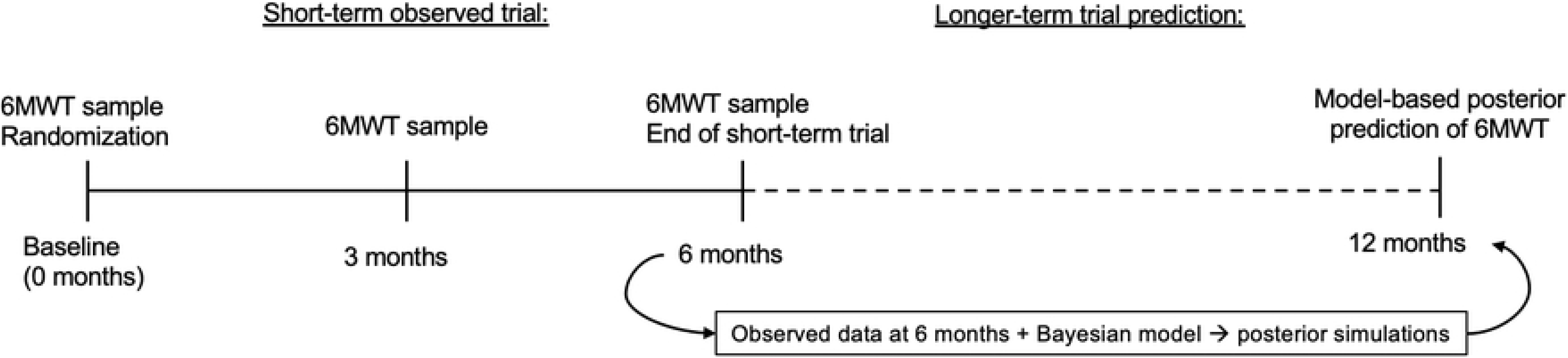
Overview of the Bayesian early decision-making framework. Individual 6-minute walk test (6MWT) samples were simulated to represent data from a short-term Duchenne muscular dystrophy (DMD) clinical trial. At completion of the trial, Bayesian model-based methods were applied to make long-term projections about the outcome at 12 months. The Bayesian model included prior natural history of the 6MWT in DMD boys.

First, replicates of short-term clinical trial data of the 6MWT were simulated from the natural history model. Data were simulated to represent the observed samples from a hypothetical short-term 6 month DMD trial. (Fig 1)

Second, the posterior probability distributions of model parameters for each trial population were estimated. The estimation was informed by prior knowledge about 6MWT natural history, along with the new data from the simulated short-term trials. Distributions of the population as well as individual parameter estimates were obtained for use in posterior simulations. (Fig 1)

Lastly, longer-term projections of the 6MWT at 12 months were made through posterior simulations. The simulations were performed conditional on the individual parameter estimates. (Fig 1)

The projected data at 12 months were analyzed under two different cohort comparison methods for decision-making: at the mean level or with individual case-matched comparisons. The short-term trial data were also analyzed using a typical frequentist method to serve as a comparator to the Bayesian approach.

These methods were applied to assess trials of various designs and three different assumed drug effect levels. The detailed methods are described in the following sections.

### Natural history Bayesian model

A Bayesian natural history model of the 6MWT in pediatric DMD patients was previously developed using full Markov chain Monte Carlo (MCMC) Bayesian methods [18]. An indirect response model with a latent process (Fig 2) was fit to literature 6MWT data with vague prior distributions [18]. Variability between subjects was quantified with an exponential random effects model, and 6MWT measurement noise was described within an additive residual error model. Refer to Lennie et. al for the full natural history model [18]. The model was validated through simulation-based methods and provided a reasonable description of the 6MWT in DMD boys [18].

**Fig 2.**
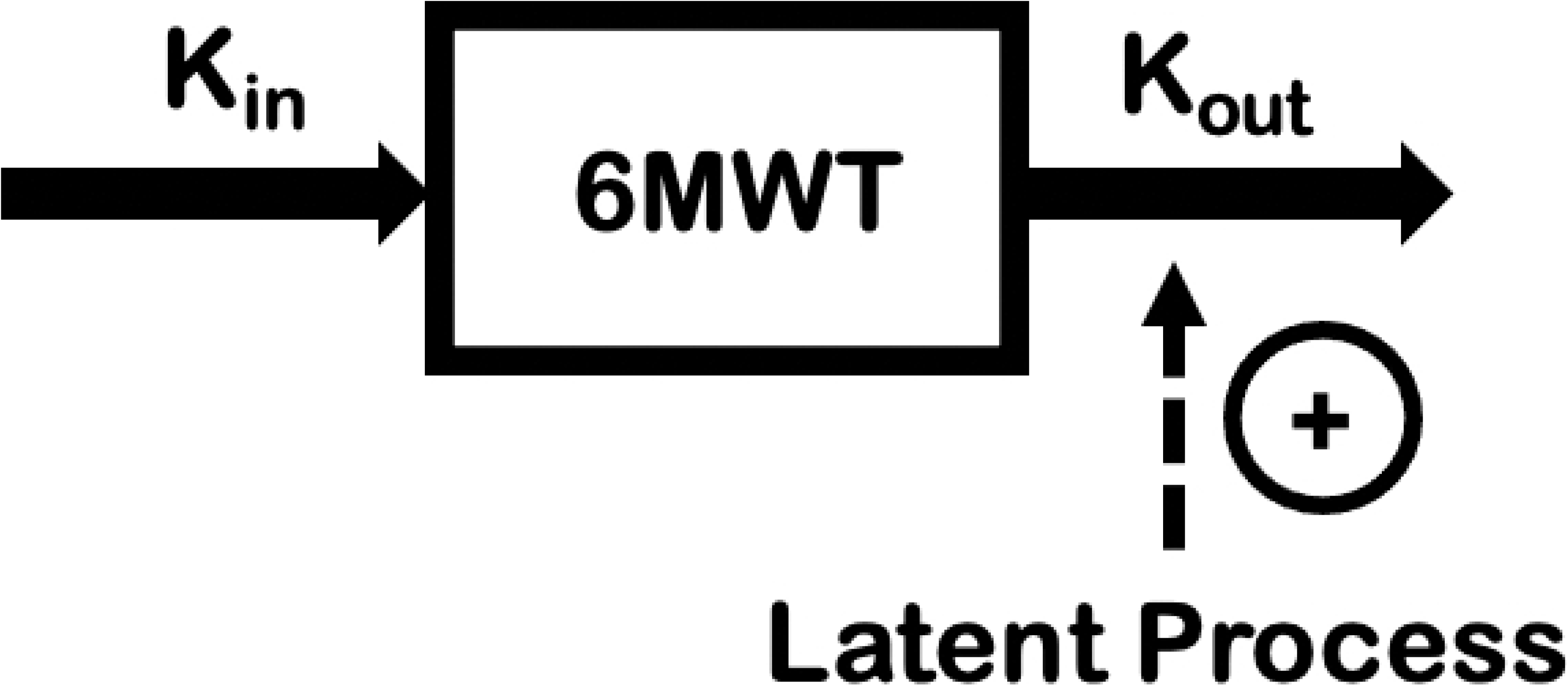
A schematic of the disease progression for DMD using the 6MWT developed by Lennie et al [18]. An indirect response model was developed simultaneously for healthy subject and DMD patient natural history data of the 6MWT. A latent process model described the effect of DMD on the 6MWT.

Individual-specific covariates were incompletely specified in the literature data [18], but a covariate analysis was performed as an exploratory investigation. The effects of corticosteroid regimen, genotype, trial type, and last sampling time were considered but none were reliably estimated given the incomplete nature of the data set. The final model did not include covariate effects.

### Hypothetical drug effect model development

A hypothetical drug effect model was developed to simulate a treatment effect on top of the 6MWT natural history trajectory. For the purpose of this paper, it was assumed that a placebo response was equal to natural history. Model structures were investigated through population mean deterministic simulations (mrgsolve v0.10.0 [19]) and calibrated to achieve three drug effect levels: 1) a low drug effect (ΔΔ6MWT <30 meters at 12 months) to represent a drug that would not meet the MCID target to demonstrate efficacy, 2) a drug effect matching the MCID of 30-meter ΔΔ6MWT at 12 months, and 3) a high drug effect (ΔΔ6MWT >30 meters at 12 months) that would demonstrate an effect of a relatively large magnitude.

### Trial simulation model development

The trial simulation model was constructed from the natural history model augmented with the hypothetical drug effect model. For purposes of trial simulation, the natural history model parameters, both fixed and random effects, were set to the means of the natural history model posterior probability distributions. The drug effect model parameters were set to the calibrated values necessary to achieve the three drug effect levels of interest.

### Simulations of short-term clinical trial data

Individual 6MWT trajectories for short-term trials were simulated with between subject variability and residual error at each of the three drug effect levels (Fig 2). The simulated trial designs reflected those of recent DMD efficacy trials [14–16], but observations were limited to 6 months in duration in order to mimic a situation where early decision making would be constrained by short term study data. A baseline age inclusion criteria of 7-13 years was applied to accurately reflect typical study populations. The prior natural history model was not informed by individual-level genotypes and corticosteroid usage data, and therefore, the simulated trials did not consider these particular inclusion criteria.

Total sample sizes and randomization schemes reflected those of typical trials (Fig 3). Patients were randomized either equally to both treatment and placebo cohorts, stratified by baseline age, or to a treatment cohort only (Fig 3). When a trial did not include a placebo arm, controls were obtained from either model-based predictions or from an external natural history database (Fig 3). The external database was created from the literature data set which was previously curated [18]. All curated 6 month samples were included in the NH database, as well as any 6 month samples that could be imputed.

**Fig 3.**
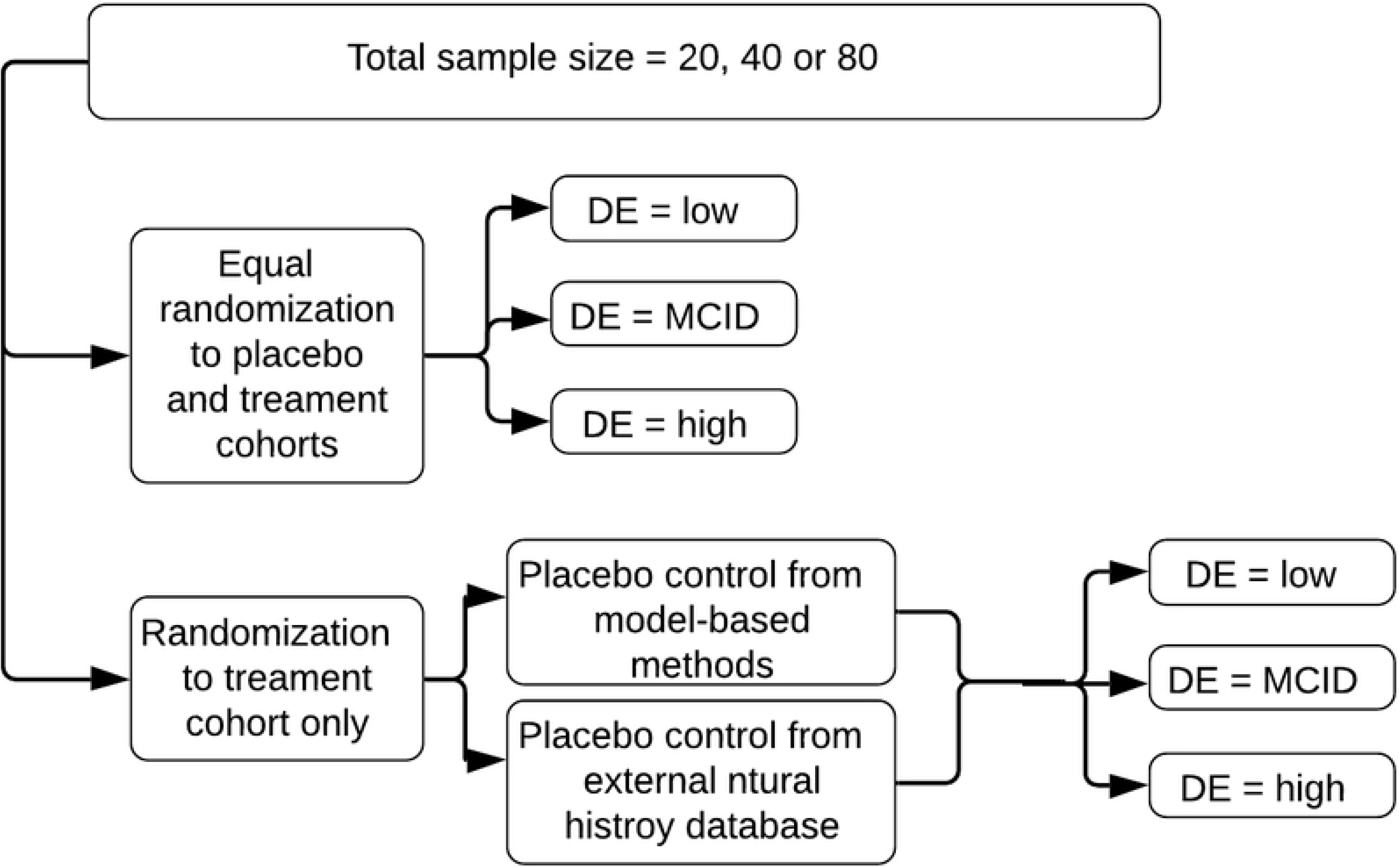
Trial design variations for the simulated short-term DMD trials. Total sample sizes varied between 20, 40 or 80 total subjects. Randomization was either equal to both placebo and treatment cohorts, or to a treatment cohort only. In the case of no placebo cohort present in the trial, controls were obtained from either a model-based prediction or an external natural history database. Drug effect (DE) sizes were defined as low, minimal clinically important difference (MCID), and high.

A number of trial simulation replicates were performed to make inferences about the population mean. Because of the extensive dimensionality of simulation scenarios and the heavy computational cost of full Bayesian modeling, 20 trial simulation replicates of each case were performed. This has been demonstrated to be a sufficient number of samples to make inferences about mean responses [20–22].

### Trial estimation model development & Bayesian estimation methods

The trial estimation model, like the simulation model, was composed of the natural history model augmented with the hypothetical drug effect model. However, the estimation model included full specification of prior distributions. The model was fit to the 6 month data from each of the 20 trial replicates for each design (Fig 3). Parameter estimation methods followed those previously described and implemented to fit the natural history model [18] (full MCMC Bayes estimation, NONMEM v7.4 [23]). Trace plots were monitored for convergence for a random selection of models from each of the trial designs.

Informative prior distributions were specified from the final parameter estimates of the natural history model. A vague prior distribution was intentionally specified for the drug effect parameter due to lack of prior information, as well as the goal to learn specifically about this component of the model given the short-term trial data.

### Posterior predictive simulations

Posterior predictive distributions of each individual’s 6MWT at 12 months were simulated from the estimated joint posterior probability distributions at each trial replicate.

The quantity of interest, the ΔΔ6MWT at 12 months, was derived from the posterior predictive distributions of the individual projections using two main cohort comparison methods: the mean treatment arm estimate, or from individual case-matching (Table 1). For individual case-matching, pairs were created based on individual-specific baseline predictors. For trials with equal randomization to both treatment and placebo cohorts, matched pairs were created from subjects in the trials. When trials only included a treatment cohort, control subjects for matching were sampled from the external natural history database. The mean of the matched pairs was summarized.

**Table 1.** Conditions of the model-based Bayesian analyses based on the implemented treatment comparison methods and decision-making methods. *Scenarios where 100 replicates of the placebo cohort source were performed.

| Cohort comparison method | Randomization scheme | Decision-making methods applied | Source of placebo reference |
| --- | --- | --- | --- |
| Difference between cohort means | Treatment & placebo | Frequentist & Bayesian model-based | Short-term trial placebo cohort |
|  | Treatment only | Frequentist & Bayesian model-based | Model-based simulated placebo reference |
| Mean of difference between individual case-matches | Treatment & placebo | Bayesian model-based | Short-term trial placebo cohort* |
|  | Treatment only | Bayesian model-based | External natural history database* |

### Go/no-go decision making

Decision criteria were applied to each trial replicate for go/no-go decision-making. Under the Bayesian paradigm, a “go” decision was declared if the posterior predictive probability of reaching the MCID at 12 months was at least 80%, otherwise a decision of “no-go” was declared. The Bayesian model-based analysis methods were applied to all trial designs (Table 1).

Frequentist analyses were applied only when observed data for both test and reference cohorts were available at the 6 month time point. This included trials with randomization to both treatment and placebo arms, or single treatment arm studies with reference control groups based on external natural history data (Table 1). Under the frequentist method, a “go” decision was declared if the 95% confidence interval (CI) lower bound from a t-test of the difference of cohort means was at least zero, otherwise a decision of “no-go” was declared. Frequentist-based decisions were also made at confidence levels of 80 and 50% to investigate the frequentist test performance at less stringent CIs.

The percentage of correct decisions under each trial design and cohort comparison method were summarized for the Bayesian and frequentist methods.

## Results

### Hypothetical drug effect model

The final hypothetical drug effect (DE) model was a logistic function on the latent process (*LP*) in the natural history model [18] (Eq 1–3). Refer to Lennie et. al [18] for the full natural history model.

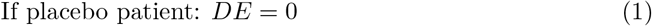

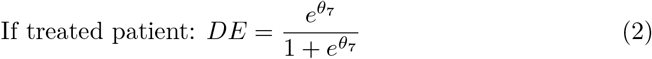

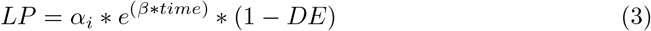

The parameter *θ*_7_ was calibrated to achieve the low, MCID, and high drug effect levels (Fig 4). The calibrated low and high drug effect levels yielded population mean ΔΔ6MWT values of 17.6 and 40 meters at 12 months, respectively. The calibrated MCID drug effect yielded the targeted 30 meter ΔΔ6MWT at 12 months. (Fig 3)

**Fig 4.**
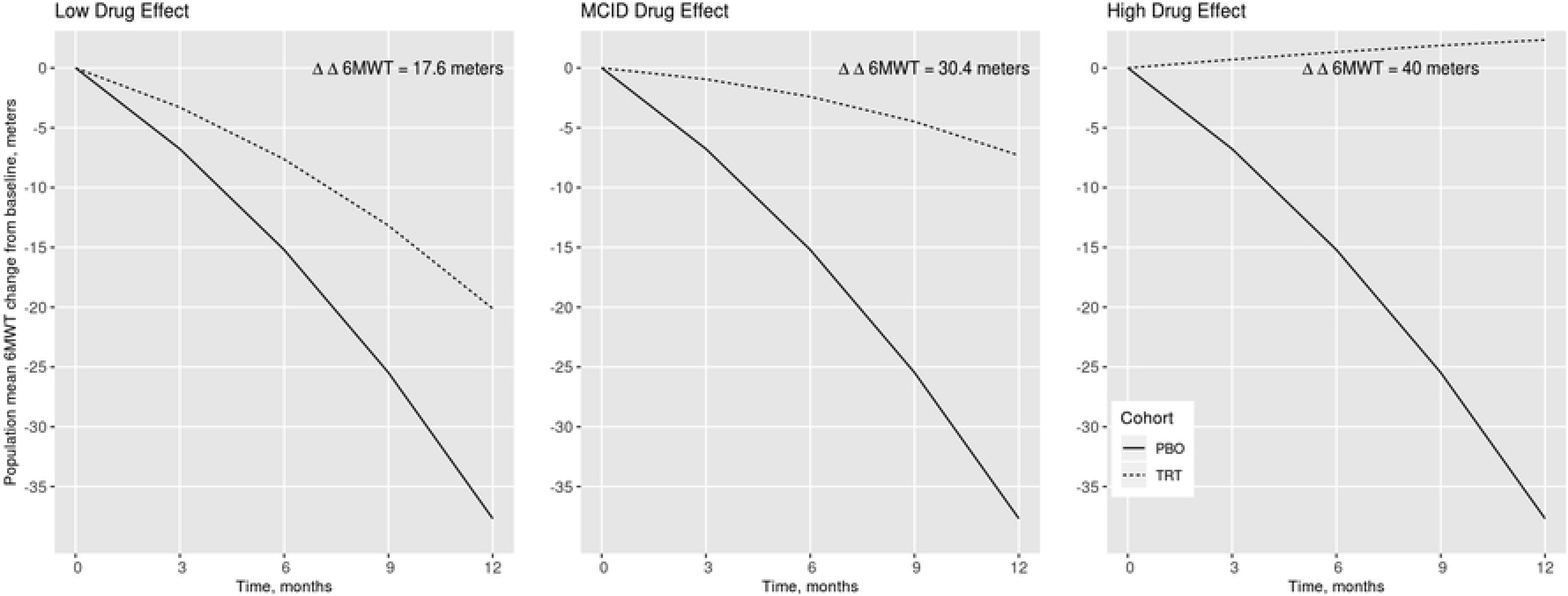
Population mean simulations of the natural history and hypothetical drug effect model at three drug effect levels (low, minimal clinically important difference (MCID), and high). The drug effect model was calibrated to achieve the three desired placebo-corrected change from baseline 6MWT (ΔΔ6MWT) values at 12 months post-baseline.

### Clinical trial simulations & posterior predictions

The trial simulation model performed as expected, with the low, MCID and high drug effects visibly discernible in the trial data from 0-6 months (Fig 5, left, points). The trial estimation model was fit to the data from each trial, with convergence confirmed by posterior parameter trace plots. The predicted individual trajectories based on observed data from 0-6 months (Fig 5, left, solid lines) were extrapolated to 12 months (Fig 5, left, dashed lines). As expected, separation between the placebo and treated posterior distributions increased with drug effect size (Fig 5, middle), as well as the probability of achieving a ΔΔ6MWT of at least 30 meters (Fig 5, right).

**Fig 5.**
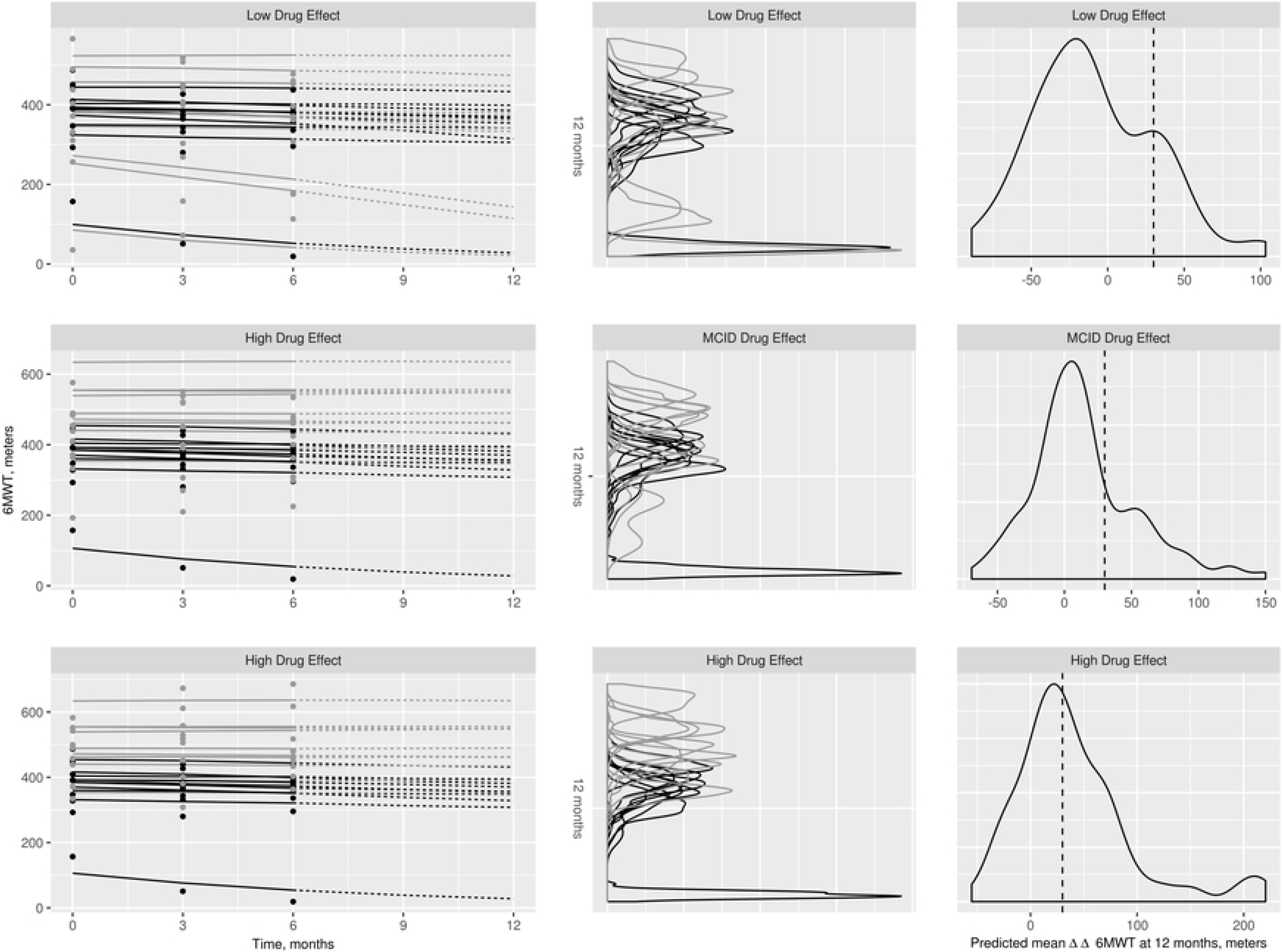
Simulated trial data, model predictions, and probability predictive distributions for one of the 20 replicates of one of the trial designs: equal randomization to both placebo and treatment arms, total sample size of 20. Left: Individual data from the short-term trials (indicated by points) with the model-predicted trajectories (solid lines indicate the short-term 0-6 month predictions, dashed lines indicate the longer-term 6-12 month predictions). The median of the posterior predictive probabilities are plotted for each time point along the solid and dashed lines. Black points and lines indicate placebo cohort, gray indicate the treatment cohort. Center: Corresponding individual posterior predictive distributions of the predicted 6-minute walk test (6MWT) at 12 months for each subject in the trials. Black lines indicate the placebo cohort, gray indicate the treatment cohort. Right: Corresponding posterior predictive distributions of the predicted mean ΔΔ6MWT at 12 months. Dashed line indicates the minimal clinically important difference (MCID) of 30 meters.

### Decision-making performance assessment

#### Bayesian model-based analysis

Performance of the Bayesian model-based methods varied across trial designs, drug effect size, and cohort comparison methods (Table 2). Overall, increased sample size resulted in improved performance. Decision-making performance was highest for the low drug effect across all trial designs and methods. Performance was moderate for the high drug effect, and poor for the MCID drug effect (Table 2).

**Table 2.** Performance of the Bayesian model-based go/no-go decision making methods across trial designs and data analysis methods. Criteria for go/no-go decisions were at least an 80% posterior predictive probability of the CFBCFP ≥30 meters at 1 year post-baseline.

|  | Cohort comparison method: mean level |  |  |  |  |  | Cohort comparison method: individual case-matched |  |  |  |  |  |
| --- | --- | --- | --- | --- | --- | --- | --- | --- | --- | --- | --- | --- |
|  | Trial design: treatment & placebo cohorts |  |  | Trial design: treatment cohort only. Placebo reference data from model-based placebo response |  |  | Trial design: treatment & placebo cohorts. 100 replications of the placebo cohort simulated from the natural history model |  |  | Trial design: treatment cohort only. Placebo reference data from 100 resampled datasets from external natural history database |  |  |
| Total N: | 80 | 40 | 20 | 80 | 40 | 20 | 80 | 40 | 20 | 80 | 40 | 20 |
| DE size: low (no-go) | 100% | 95% | 90% | 100% | 100% | 100% | 100% | 100% | 100% | 100% | 90% | 85% |
| DE size: MCID (go) | 0% | 5% | 10% | 0% | 5% | 5% | 0% | 0% | 5% | 25% | 25% | 30% |
| DE size: high (go) | 25% | 15% | 10% | 15% | 15% | 20% | 17% | 20% | 15% | 100% | 70% | 55% |

Inconsistencies were evident in the case-matched method using the external database controls compared to the other three cohort comparison methods. The external database method performed markedly better than the others for the MCID and high drug effects. Performance ranged from 25-30 and 55-100% for the MCID and high drug effects under the external database case, but only 0-10 and 15-25% for the other three cases, respectively.

#### Frequentist analysis

Performance using the frequentist analysis was poor across all trial designs and cohort comparison methods. The average decision accuracy rate across all scenarios was 8.98%. Only four cases made correct go/no-go decisions in at least 50% of the replicates, with three of those four cases for the low drug effect. Performance was highest at the lowest CI level (50% CI). There was no clear pattern of increased performance with increased sample size or increased drug effect. (Table 3)

**Table 3.** Performance of the frequentist analysis go/no-go decisions across trial designs and analysis methods. Go/no-go decisions were made at frequentist confidence levels of 95, 80 and 50%.

| Go/no-go decision criteria: Go if the LB ≥ 0 for a CI of: | Cohort comparison method: mean level |  |  |  |  |  |  |  |  |  |  |  |  |  |  |  |  |  |
| --- | --- | --- | --- | --- | --- | --- | --- | --- | --- | --- | --- | --- | --- | --- | --- | --- | --- | --- |
|  | Trial design: treatment & placebo cohorts |  |  |  |  |  |  |  |  | Trial design: treatment cohort only. Placebo reference data from external natural history database |  |  |  |  |  |  |  |  |
|  | 95% |  |  | 80% |  |  | 50% |  |  | 95% |  |  | 80% |  |  | 50% |  |  |
| Total N: | 80 | 40 | 20 | 80 | 40 | 20 | 80 | 40 | 20 | 80 | 40 | 20 | 80 | 40 | 20 | 80 | 40 | 20 |
| Truth: No-go | 5% | 0% | 10% | 15% | 0% | 35% | 25% | 0% | 65% | 0% | 15% | 5% | 0% | 40% | 0% | 85% | 65% | 20% |
| Truth: Go, MCID DE | 0% | 0% | 5% | 0% | 0% | 15% | 0% | 0% | 55% | 0% | 0% | 0% | 0% | 0% | 0% | 0% | 5% | 5% |
| Truth: Go, High DE | 0% | 0% | 0% | 0% | 0% | 5% | 0% | 0% | 10% | 0% | 0% | 0% | 0% | 0% | 0% | 0% | 0% | 0% |

## Discussion

This paper presented an early decision-making framework for rare disease drug development. The framework was applied to hypothetical short-term DMD trials of varying designs to make go/no-go decisions about continuation to late stage trials. Decisions were informed by prior natural history combined with newly collected trial data, and based on long-term projections from Bayesian model-based methods. The probability of achieving the targeted demonstration of efficacy was determined for 20 replicates of each trial design. Frequentist tests were also applied to the 20 replicates of each trial design, determining the probability of a difference between the treatment and placebo cohort means. Performance across the 20 replicates was summarized. The proposed Bayesian model-based framework was superior to the frequentist method for making go/no-go decisions across all trial designs and analysis methods in DMD. The frequentist control method was limited to short-term trial data only, while the Bayesian methods were powered with both observed and prior information.

As demonstrated, there were inconsistencies evident between the results using the observed external database versus model-based placebo control data. Performance was markedly improved when case-matched control subjects were obtained from the external database (Table 2). The external database consisted of observed data curated from the literature, while all other cases used model-simulated control data.

Investigations were performed to identify the possible source of the inconsistency between the observed external database and the model-simulated placebo control data. The results of a simulation-based evaluation of the natural history model indicated bias in model predictions that may have propagated into the model-simulated control data. Simulation-based evaluations of the 6MWT change from baseline (CFB) indicated positive bias in the tails of the model predicted CFB distributions (Fig6). Positive bias would cause some simulated placebo responses to be less severe than those of the observed data, making the drug effect appear smaller. Decreases in the apparent drug effect would cause error in decisions for the MCID and high DE cases, but would still lead to a “no-go” decision for the low drug effect level. This hypothesis is in accordance with the results: performance was similar across all 4 analysis methods for the low drug effect, but was markedly higher for MCID and high drug effects using the external database controls.

**Fig 6.**
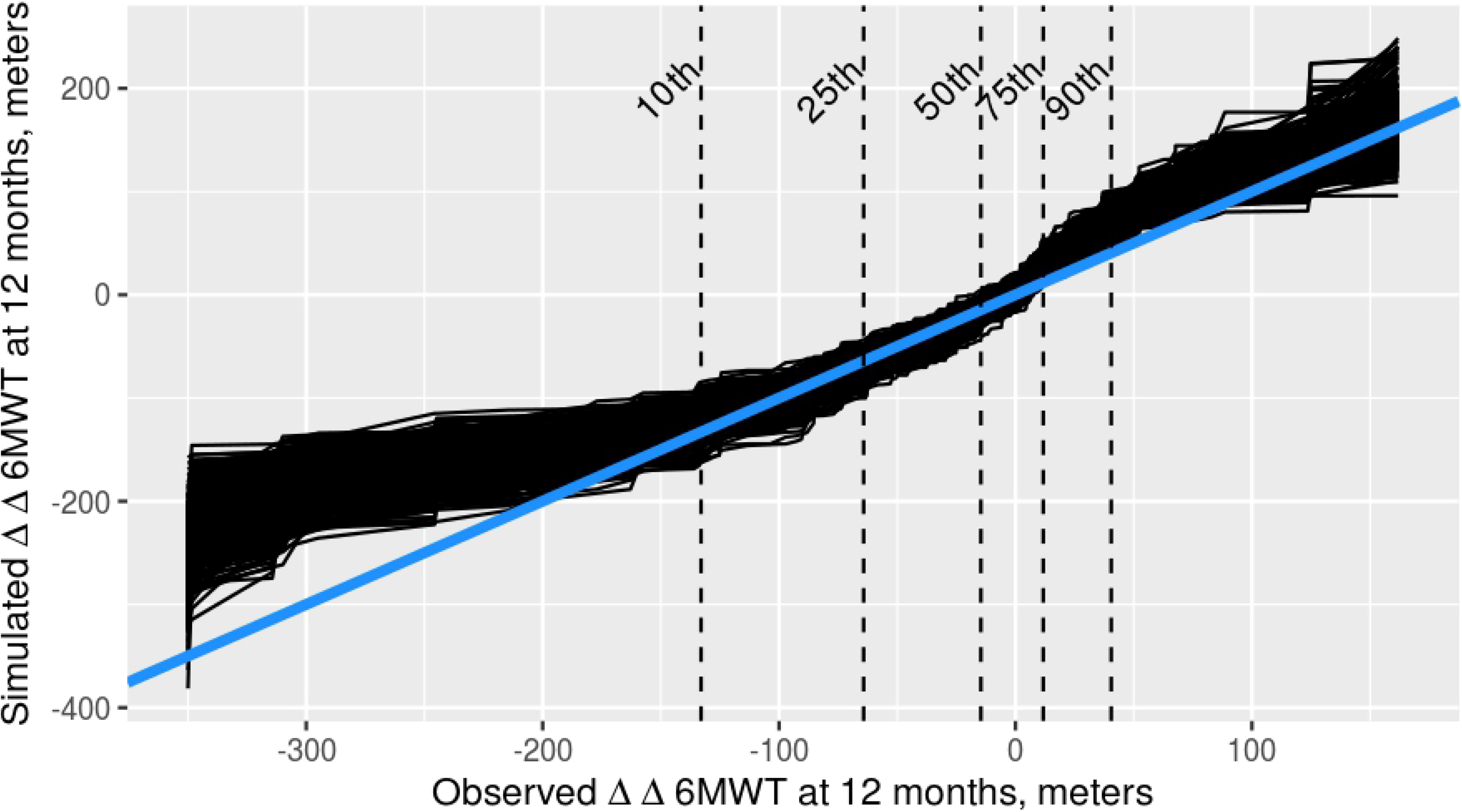
Predictive check quantile-quantile plot for the natural history model fit to a subset of subjects with baseline ages between 7-13 years. The simulated 6-minute walk test (6MWT) change from baseline (CFB) is plotted against the observed 6MWT CFB from 500 Monte Carlo simulations. The 10th, 25th, 50th, 75th and 90th quantiles of the observed 6MWT CFB distribution are shown with dashed lines. The blue line indicates the line of identity.

The model evaluation results on the CFB scale indicated that a predictor(s) to describe the relationship between baseline and post-baseline points may be missing. Though the natural history model was previously validated to predict the 6MWT, it had not been previously evaluated specifically on the CFB scale in the age 7-13 subset of patients [18]. The example using DMD was limited to the literature data, and covariate models were not supported. When lacking covariates, optimization methods such as the iterative back step method [**?**] could be applied to train the model to the dataset, but the ideal simulation model would include individual-specific covariates. Ideally, complete individual-level data should be collected and the model should be well validated for its specific use in a drug development program.

Due to issues with model misspecification on the 6MWT CFB scale, inferences about optimal methods for DMD early go/no-go decisions should be viewed as preliminary. Results for the cases using model-simulated placebo controls may be subject to model bias, while those using observed external data controls are likely to be reliable. Decision-making performance was strong for the low and high drug effects when using the external database, and increased with sample size (Table 2). At the MCID drug effect level, performance was moderate (25-30%, Table 2). These results indicate that drug effects with larger magnitudes, whether positive or negative, were well predicted, but accurate decision-making for a minimal drug effect is difficult. These results align with characteristics of the 6MWT endpoint: the endpoint is highly variable between patients, so the effects of the MCID drug may be confounded by 6MWT variability, while drug effects of high magnitudes may be more detectable. The MCID drug effect may be more readily detected with larger sample sizes or in a longer study. These results may be considered when designing confirmatory trials in DMD with the 6MWT endpoint; demonstration of efficacy at 12 months for a drug whose true population effect is close to the 30 meter ΔΔ6MWT is likely to be challenging given typical DMD sample size constraints.

The specific results of this DMD case study should be reevaluated using a full dataset complete with individual predictors. These limitations aside, the framework presented here provides a proof of concept for the utility of Bayesian model-based methods for decision making in rare disease trials.

The small sample sizes, lack of standardized outcome endpoints, and limited clinical trial data inherent in rare diseases create a unique need for open science collaborations and pre-competitive data sharing. The results of the DMD example highlight the challenges of validating a model for its intended use without access to a complete individual level data set. Only five published DMD studies reported the 6MWT data at the individual level [9, 24–27] and most studies reported natural history data in summary form only. Open access databases of individual-level patient data would contribute greatly to the understanding of rare diseases by facilitating data-driven and model informed trial designs and drug development decision making.

